# Dual mRNA delivery corrects physical and biochemical deficits in a mouse model of trifunctional protein deficiency

**DOI:** 10.64898/2026.11.19.767734

**Authors:** Troy von Beck, Tatiana Terranova, Bahjat Fadi Marayati, Haoyue Zhang, Liyang Zhu, Songtao Li, Bruce R. Donald, Sarah P. Young, Myles Lindsay, Christopher A. Todd, Dwight D. Koeberl

**Affiliations:** Division of Medical Genetics, Department of Pediatrics, Duke University Medical Center, Durham, NC, USA; Duke Human Vaccine Institute, Duke University School of Medicine, Durham, NC, USA; Duke Medicine Biochemical Genetics Laboratory, Duke University, Durham, NC, USA; Department of Computer Science, Duke University, Durham, NC, USA; Department of Computer Science, Department of Chemistry, Department of Mathematics, Duke University, Department of Biochemistry, Duke University School of Medicine, Durham, NC, USA

**Keywords:** Trifunctional protein deficiency, Lipid nanoparticle, Therapeutic mRNA, Fatty acid oxidation, Dominant negative, Rare disease, Mouse model

## Abstract

Mitochondrial trifunctional protein deficiency (TFP-D) is a rare, life threatening long-chain fatty acid oxidation disorder caused by variants in either the HADHA or HADHB subunits, leading to toxic metabolite accumulation and systemic energy deficiency. Current dietary management strategies are only partially effective and fail to prevent long term morbidity and mortality. Here, we developed a lipid nanoparticle (LNP) mediated mRNA replacement strategy to restore TFP function in a moderate severity TFP-D mouse model by delivering both Hadha and Hadhb mRNA. In cultured TFP-D mouse fibroblasts, dual mRNA delivery, but not single subunit replacement, rescued long-chain fatty acid oxidation and revealed a dominant negative effect exerted by the endogenous M404K variant Hadhb. *In vivo*, intravenous administration of dual mRNA-LNPs restored hepatic TFP expression, reduced circulating long-chain acylcarnitines, and improved physiological performance in aged TFP-D mice. Notably, intravenous delivery of dual mRNA LNPs significantly rescued cold tolerance test (CTT) performance by up to 71.6% compared to untreated and healthy controls. Together, these findings establish dual subunit LNP-mRNA delivery as a promising therapeutic strategy for TFP-D and support its potential applicability to other long-chain fatty acid oxidation disorders.

## Introduction

Mitochondrial trifunctional protein (TFP) is a catabolizing enzyme composed of two alpha (HADHA) and two beta (HADHB) subunits, which together mediate 3 of the 4 steps in long-chain fatty acid oxidation. In the β-oxidation spiral, the sequential steps of long-chain enoyl-CoA hydration and 3-hydroxyacyl-CoA oxidation are mediated by HADHA, while the final step of β-ketoacyl-CoA thiolysis is performed by HADHB. Variants in either *HADHA* or *HADHB* that disrupt the whole enzyme complex are causative for the associated TFP deficiency disorder (TFP-D), while isolated loss of 3-hydroxyacyl-CoA oxidation by variants in *HADHA* drive the related, but less severe long-chain 3-hydroxyacyl-CoA dehydrogenase deficiency (LCHAD-D).^1, 2^ Cells with reduced or absent TFP have impaired β-oxidation of long chain fatty acids (14-20 carbon atoms) with less significant impairment for medium chain fatty acids (8-12 carbon atoms), owing to redundant processes.^3^ Consequently, cells deficient for the enzyme complex are unable to generate acetyl-CoA from long-chain fatty acids for energy production and accumulate cytotoxic long-chain metabolites.^4^

Collectively, the incidence of fatty acid oxidation disorders is between 1:5,000-1:10,000 births, while the incidences for TFP-D and LCHAD-D individually are estimated to be 1:750,000 and 1:250,000 respectively.^5, 6^ In TFP-D patients, the combined cytotoxicity and metabolic insufficiency manifest systemically, with common symptoms of hypoketotic hypoglycemia, cardiomyopathy, encephalopathy, liver dysfunction, episodic rhabdomyolysis, hypotonia, peripheral neuropathy, and retinopathy.^1, 2^ The severity of these symptoms correlates with the degree of TFP enzyme dysfunction, as complete loss of enzyme activity causes the lethal neonatal form with cardiomyopathy, while partial enzyme function results in the early-onset hepatic form or milder late-onset neuromyopathic form.^7^

Current strategies for disease management are limited to lifestyle and diet changes, with avoidance of fasting and prolonged exercise combined with a high carbohydrate, low fat diet supplemented with medium chain triglycerides or triheptanoin.^1, 8^ While these interventions can delay symptom onset and progression, patients remain at elevated risk of mortality, long-term complications, and episodes of acute metabolic decompensation from stressors such as infection.^1, 9^ Therefore, new therapeutic interventions are warranted to improve patient quality of life and progression-free survival. With recent advances in lipid nanoparticle (LNP) formulations and synthesis of modified RNA molecules, mRNA has become an attractive vector for the delivery of protein replacement therapies. Indeed, two recent studies produced encouraging results using LNP-mRNA complexes to biochemically and physiologically correct fatty acid oxidation disorders both *in vitro* and *in vivo.*^10, 11^

To date, two mouse models of TFP-D have been developed, which present either the severe neonatal phenotype from a homozygous Hadha knockout or the early-onset hepatic phenotype from a homozygous Hadhb M404K missense variant.^12, 13^ In this study, we utilized the early-onset hepatic mouse model to develop and characterize an LNP-mRNA therapeutic for protein replacement therapy. We further identified a substantial dominant negative effect from the endogenous *Hadhb* missense variant, which could be overcome by delivering both the Hadha and Hadhb subunits as mRNA. Delivery of LNP-Hadha/Hadhb mRNA *in vivo* significantly improved metabolic performance, supporting therapeutic LNP-mRNA as a viable option for TFP-D treatment.

## Results

### Dual Hadha/Hadhb mRNA transfection rescues long-chain fatty acid oxidation in vitro

To establish an *in vitro* TFP-D model for testing mRNA therapies, skin fibroblasts were isolated from wild-type (WT) or Hadhb^M404K/M404K^ (TFP-D) mice and grown in a minimal medium supplemented with BSA-conjugated palmitic acid (C16:0) and L-carnitine. As the TFP-D fibroblasts cannot efficiently metabolize long-chain fatty acids via β-oxidation, there is an accumulation of long-chain acylcarnitine intermediates in the culture medium which can be detected by liquid chromatography tandem mass spectrometry (LC-MS/MS) method (Figure 1A-C). TFP-D fibroblasts demonstrated statistically significant increases in C16:0 and C16:0 derivative LCFA species. Curiously, TFP-D fibroblasts also produced increased concentrations of medium-chain acylcarnitine intermediates.

**Figure 1.**
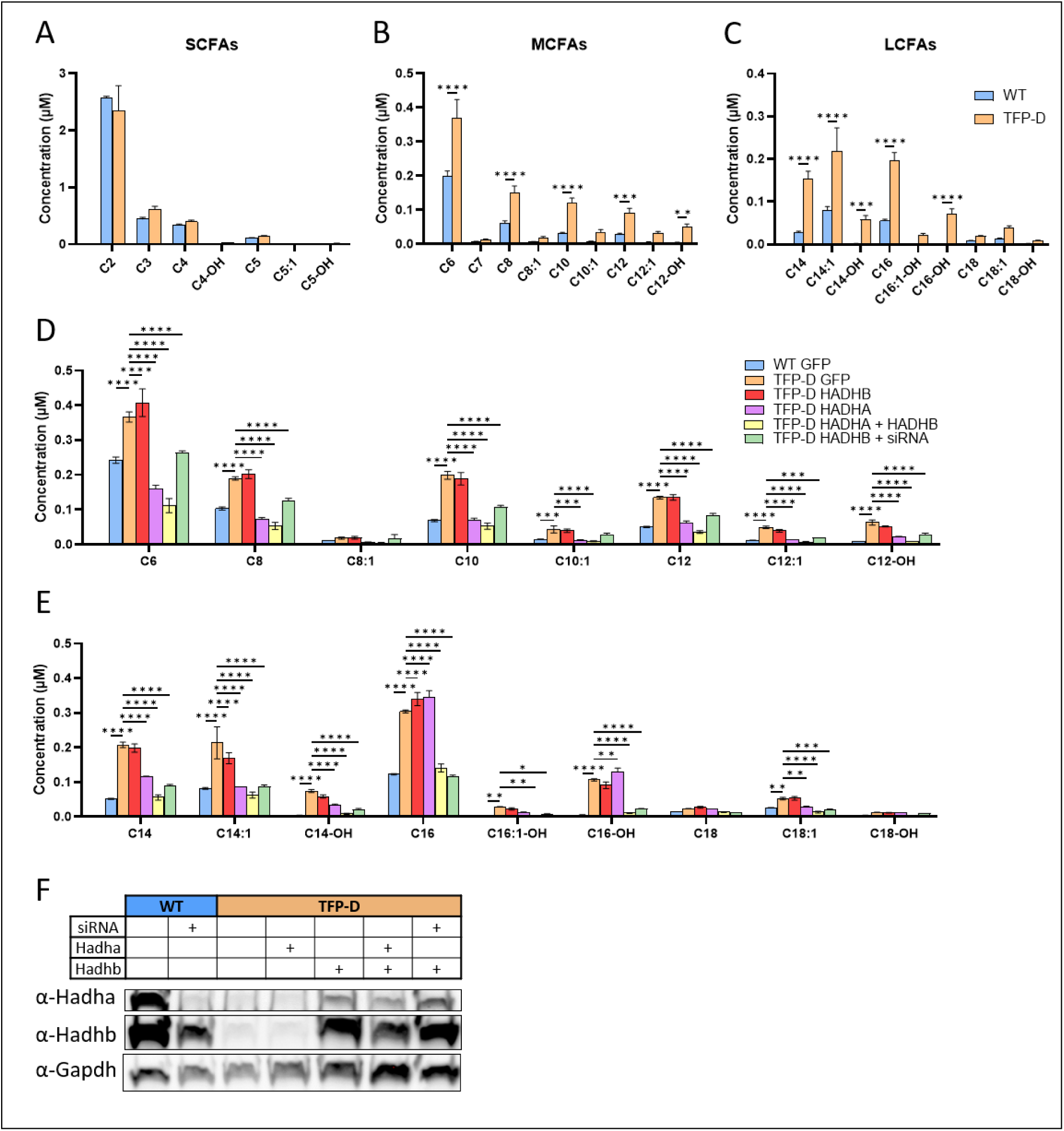
Accumulation of long and medium chain acylcarnitines is alleviated in TFP-D fibroblasts following mRNA replacement therapy. (A-C) Medium and long chain acylcarnitines are increased in the culture medium TFP-D fibroblasts following a C16:0 challenge. WT or TFP-D fibroblasts were cultured in low glucose MEM supplemented with 200µM C16:0. The challenge medium was collected after a 72-hour incubation and the acylcarnitine intermediate concentrations determined by LC-MS/MS. Data are presented as mean ± standard deviation from three independent experiments (n = 3). Two-way ANOVA with Šidák multiple comparisons test. (D-E) Medium-chain and long-chain acylcarnitines in challenge medium 72 hours following mRNA/*Hadhb*-siRNA transfection. Single transfections with *GFP*, *Hadha*, or *Hadhb* used 2µg of mRNA, while double transfections used 1µg of each component for the same total RNA dose. Data are presented as mean ± standard deviation from three independent experiments (n = 3). Two-way ANOVA with Dunnet’s multiple comparisons test. (F) Representative blot of Hadha and Hadhb protein expression in untreated control fibroblasts and fibroblasts collected on day 3 after transfection. *p<0.05, **p<0.01, ***p<0.001, ****p<0.0001

To investigate the ability of mRNA replacement to correct the FAO defect, TFP-D fibroblasts were transfected with *in vitro* transcribed (IVT) Hadhb and/or Hadha mRNA and compared to either WT or TFP-D fibroblasts receiving a GFP control mRNA (Figure 1D-E and Figure S2). It was observed that transfection of either WT or TFP-D fibroblasts with GFP control mRNA increased the concentration of medium-chain and long-chain acylcarnitines relative to untreated cells, which may reflect toxicity of the lipofectamine or an innate immune response to the mRNA cargo. Notably, cells treated with an equivalent amount of *Hadhb*mRNA did not substantially recover and only produced marginal improvements in C2 and C14:1 concentrations.

It has been previously reported that restoration of long-chain 3-ketoacyl-CoA activity in human TFP-D fibroblasts required delivery of plasmids encoding both the HADHA and HADHB subunits, even when the patient variants were limited to HADHB.^14^ To evaluate this possibility, TFP-D fibroblasts were transfected with either Hadha mRNA alone or a 1:1 (w/w) mixture of Hadha/Hadhb mRNA and compared to the previous transfection controls. Treatment by combined delivery of both *Hadha* and *Hadhb* mRNA significantly improved concentrations of all medium-chain and long-chain acylcarnitines in TFP-D fibroblasts, essentially restoring them to WT levels. Unexpectedly, transfection with *Hadha* mRNA alone also corrected the accumulation of medium-chain acylcarnitines, but did not improve C16 and only moderately improved C14 species (Figure 1D-E).

Based on these findings, it was suspected that the endogenous M404K variant Hadhb subunits limits the cellular response to replacement *Hadhb* mRNA transfection by competitively binding to endogenous Hadha subunits in a “dominant negative-like” manner, which is only overcome when Hadha is also supplied in excess. Therefore, a follow-up experiment was conducted in which WT *Hadhb* mRNA was co-transfected with a *Hadhb* siRNA that binds to the 3’UTR of Hadhb and selectively depletes the endogenous transcript by up-to 80% (Figure S1). When combined, the depletion of endogenous M404K variant Hadhb subunits and repletion with WT Hadhb effectively rescued the TFP-D phenotype similar to the co-transfection of *Hadha* and *Hadhb* mRNA (Figure 1D-E). This strongly supports the existence of dominant negative Hadhb variant activity, which must be overcome to achieve an effective clinical response.

Paradoxically, transfection of TFP fibroblasts with *Hadha*or *Hadhb* mRNA alone produces inverse results when quantifying disease correction by either western blot (Figure 1F) or acylcarnitine profiling (Figure 1D-E). While treatment with *Hadha* mRNA alone improved medium-chain and some long-chain acylcarnitines, it did not increase the expression of *Hadha* or *Hadhb*. Conversely, treatment with *Hadhb* mRNA alone did increase the expression of both *Hadha* and *Hadhb*, but failed to improve any medium-chain or long-chain acylcarnitines. This discordance was stable across multiple assays and two separate mRNA preparations.

### Endogenous TFP enzyme activity is limited by Hadha expression

Given the unexpected normalization of medium-chain acylcarnitine intermediates in fibroblasts transfected with *Hadha* mRNA alone, it was hypothesized that the endogenous expression of *Hadha* may be a rate limiting factor in the assembly of trifunctional protein oligomers. Even though *Hadha* and *Hadhb* share many of the same expression determinants due to their uniquely shared bidirectional promoter and head-to-head orientation, it was theorized they may have discordant expression due to either differential transcription in the sense and antisense directions, differential stability of the encoded mRNAs, or differential rates of protein translation.

In support of this theory, absolute quantification of the *Hadha* and *Hadhb* mRNA transcripts in WT and TFP-D mouse fibroblasts by RT-qPCR demonstrated a 2.85-fold higher copy number of Hadhb transcripts relative to Hadha (Figure 2A). Further, there is no significant difference in the expression of either transcript between WT and TFP-D fibroblasts, suggesting transcription and mRNA stability is not affected by the M404K variant. This analysis was additionally extended to characterization in human tissues using the Human Protein Atlas (HPA) and Genotype-Tissue Expression (GTEx) mRNA-sequencing datasets. While the HPA dataset demonstrated a similar pattern of excess HADHB transcripts across 4 key tissues, this pattern was less pronounced and more variable in the GTEx dataset (Figure 2B).

**Figure 2.**
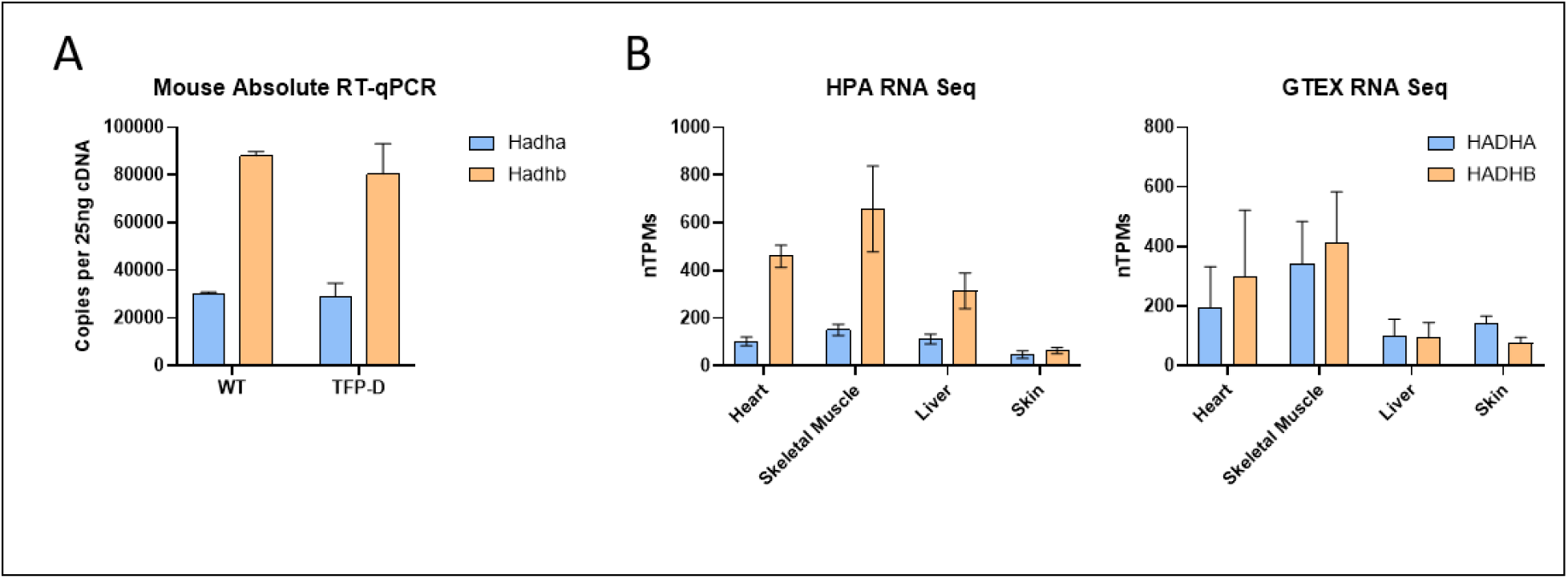
Expression of endogenous trifunctional protein subunits in mouse fibroblasts and human tissues. (A) Absolute RT-qPCR quantification of *Hadha* and *Hadhb* transcripts in untreated, cultured fibroblasts from WT and TFP-D mice. Data are presented as mean ± standard deviation from three independent experiments (n = 3). (B) Normalized abundance of *HADHA* and *HADHB* transcripts across 4 tissues in the Human Protein Atlas (HPA) and Genotype-Tissue Expression (GTEx) RNA sequencing datasets.

**Figure 3.**
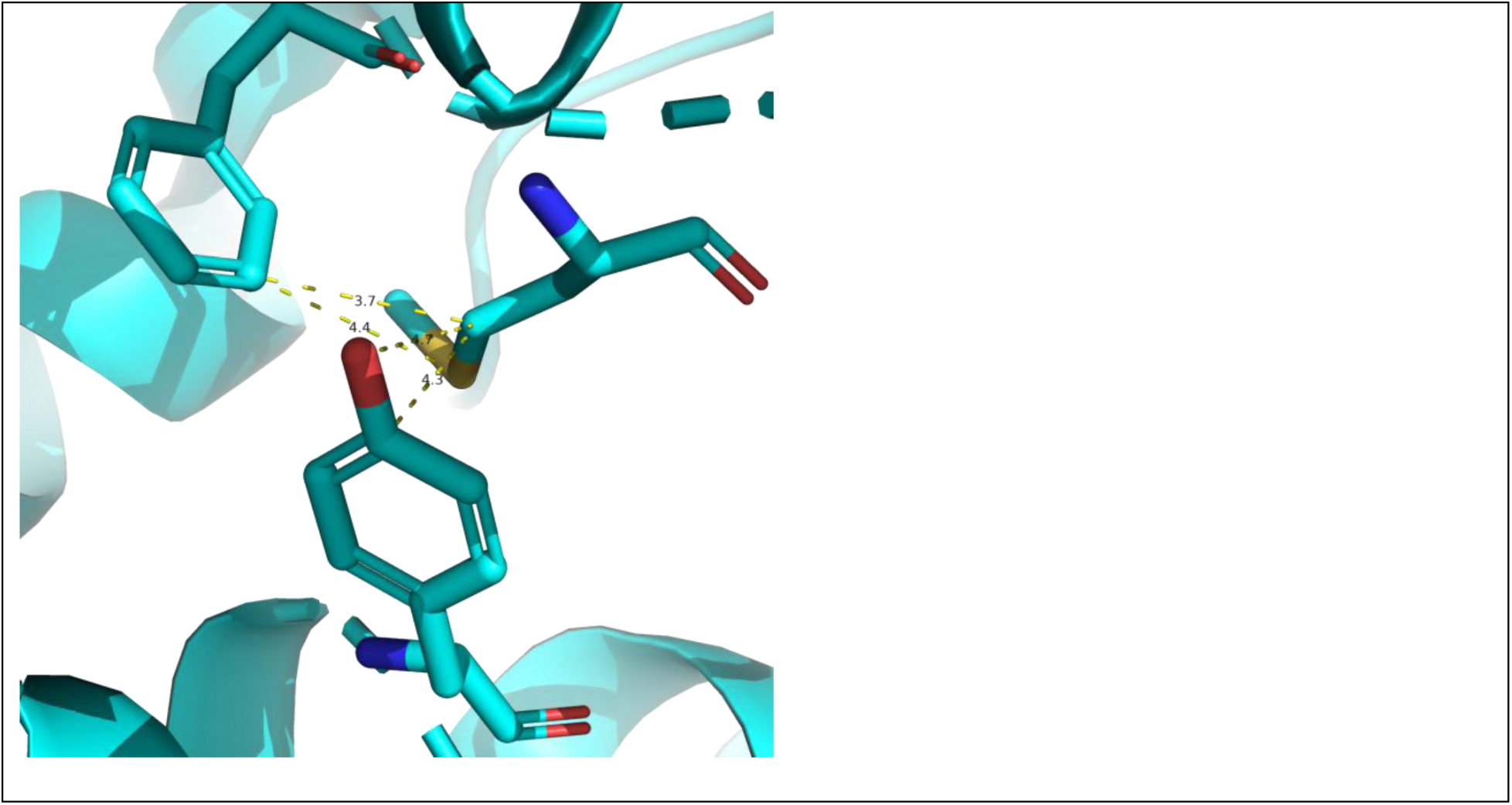
Structural model of the Met404 Aro–Met–Aro motif in Hadhb (PDB: 6DV2, chain B). Met404 (sticks) is flanked by Y358 and F399 (sticks). Geometric analysis supports a well-defined sulfur–aromatic bridge consistent with previously reported interaction geometries.^17, 18^ The Met404–F399 contact (Sδ···centroid distance = 5.49 Å; polar angle θ = 89.9°, defined relative to the aromatic ring normal) adopts an edge-on geometry, with the sulfur positioned approximately within the plane of the phenyl ring near CD1 (Sδ···CD1 = 4.34 Å). In contrast, the Met404–Y358 contact (d = 5.20 Å; θ = 18.1°) adopts a face-on geometry, with the sulfur positioned above the tyrosine ring plane (S-to-plane distance = 4.94 Å). Both interactions satisfy the commonly used d ≤ 6 Å sulfur–aromatic contact criterion, supporting assignment of a single methionine Sδ simultaneously engaging two aromatic π systems.

### Computational ensemble-based thermodynamic modeling reveals the M404K variant disrupts a stabilizing methionine-aromatic bridge

To quantitatively assess the impact of the M404K variant on Hadhb protein stability, analysis of computational protein stability was performed using the PAStE (Predicting the Affect on Stability using Ensembles) algorithm implemented in the OSPREY protein design software suite.^15^ Starting from the crystal structure of the human mitochondrial trifunctional protein (PDB: 6DV2), substitution of Met404 (Met403 in PDB 6DV2) with lysine was modeled. As the high concentration of acyl-CoA species in the mitochondria causes spontaneous acylation of lysine residues,^16^ two common post-translationally modified lysine derivatives Nε-acetyl-lysine (ALY) and Nε-succinyl-lysine (SLL) were also modeled to evaluate how such modifications would affect protein stability.

Initial inspection of the Hadhb subunit identified a conserved Aro–Met–Aro motif involving Y358, M404, and F399. Sulfur–aromatic interactions are non-covalent contacts in which the thioether sulfur (Sδ) of methionine interacts favorably with the π system of an aromatic residue (Phe, Tyr, or Trp), typically contributing ∼1–3 kcal mol⁻¹ of stabilization beyond nonspecific hydrophobic packing. Such interactions are frequently observed in protein structures and have been implicated in both structural stabilization and functional regulation. Using geometric criteria adapted from prior studies,^17, 18^ a Met-aromatic contact was defined when the distance between the methionine Sδ atom and the aromatic ring centroid was ≤ 6 Å. The interaction geometry was further characterized using the angle θ between the Sδ → centroid vector and the aromatic ring normal, where smaller θ values correspond to more face-on arrangements and larger θ values to more edge-on geometries. An Aro–Met–Aro bridge was assigned when a single methionine Sδ simultaneously satisfied these criteria with two aromatic partners. To model the effect of each lysine variant, the conformational space was defined to include continuous rotameric flexibility at the mutation site (residue 404) and eight surrounding residues: T357, Y358, A359, W398, F399, A400, Y403, G405. Notably, three of these flexible residues Y358, F399, and Y403 form the aromatic cage surrounding M404, including the Y358–M404–F399 methionine–aromatic bridge that is central to this study. PAStE computed provably accurate ΔΔG values (wild-type to mutant free energy differences) with guaranteed error bounds (Table 1). Both the unmodified lysine and acylated lysine variants at position 404 were predicted to decrease protein stability relative to the wild-type methionine, with the succinyl-lysine modification (ΔΔG = 3.14 kcal/mol) exerting a greater destabilizing effect than acetyl-lysine (ΔΔG = 2.27 kcal/mol) or unmodified lysine (ΔΔG = 2.07 kcal/mol). This is consistent with the larger steric bulk of the succinyl group, which likely exacerbates steric strain within the methionine–aromatic bridge (Figure S3). The positive ΔΔG values indicate that post-translational acylation of the M404K variant does not compensate for the loss of native methionine–aromatic interactions and instead further compromises structural stability.

**Table 1.**
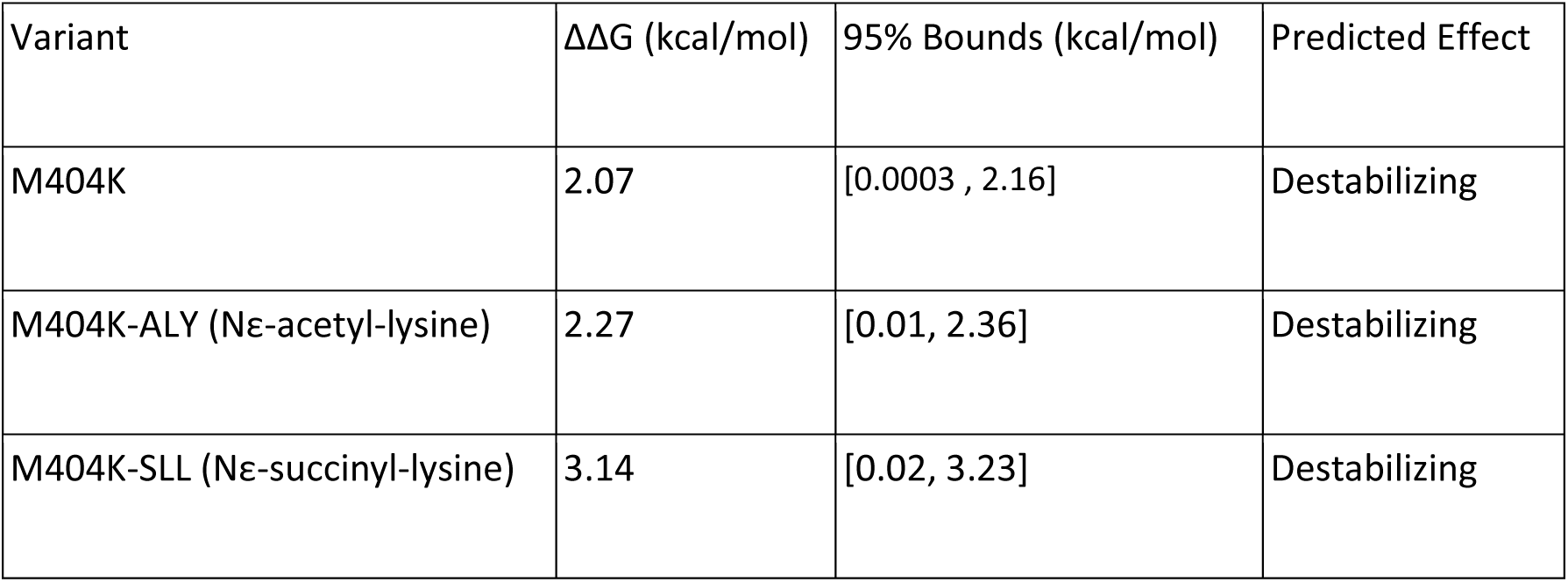
PAStE-computed ΔΔG values for post-translational modification variants at position 404 in Hadhb.

| Variant | ΔΔG (kcal/mol) | 95% Bounds (kcal/mol) | Predicted Effect |
|---|---|---|---|
| M404K | 2.07 | [0.0003 , 2.16] | Destabilizing |
| M404K-ALY (Nε-acetyl-lysine) | 2.27 | [0.01, 2.36] | Destabilizing |
| M404K-SLL (Nε-succinyl-lysine) | 3.14 | [0.02, 3.23] | Destabilizing |

These computational results corroborate the experimental observation that the M404K variant disrupts complex stability and provide additional evidence that post-translational modifications at this position do not rescue the stability defect.

### Subcellular localization and expression kinetics of Hadha/Hadhb following in vitro transfection

To confirm that transfected *Hadha* and *Hadhb* mRNA achieve normal cellular expression and mitochondrial localization, transfected fibroblasts were characterized by immunofluorescence microscopy (Figure 4A). In WT fibroblasts, both Hadha and Hadhb subunits localize to mitochondria as indicated by their characteristic “dashed web” staining pattern within cells. This pattern was more easily visualized for Hadha than Hadhb, owing to differences in the sensitivity of the primary antibodies. In TFP-D fibroblasts, the staining intensity for both Hadha and Hadhb was significantly reduced. As expected, based on the previous results, transfection of cells with both *Hadha* and *Hadhb* mRNA resulted in a normal staining pattern for each subunit.

**Figure 4.**
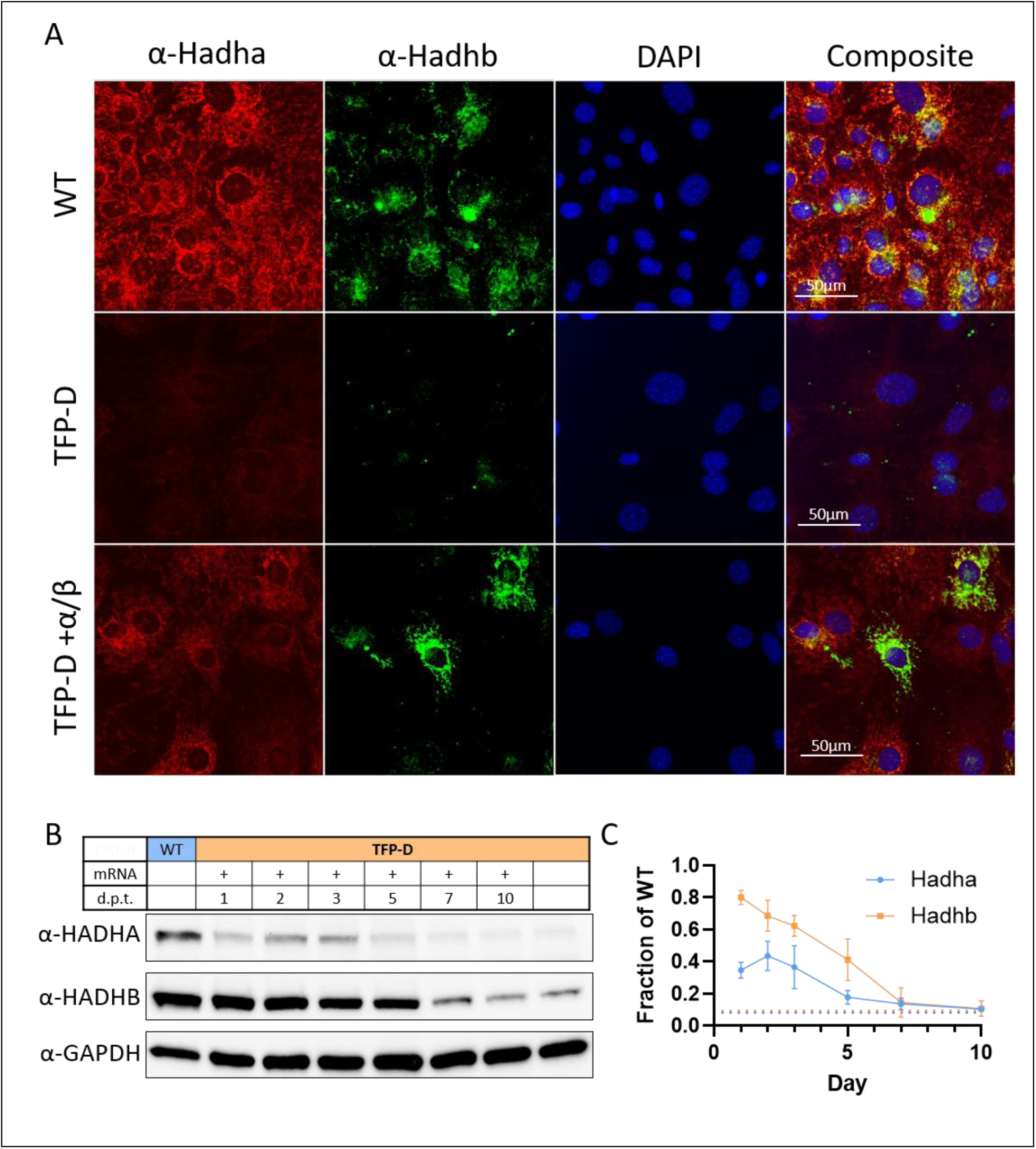
Subcellular localization and expression kinetics of Hadha and Hadhb from IVT mRNA. (A) TFP-D fibroblasts transfected with *Hadha* and *Hadhb* mRNA show normal protein subcellular localization. (B) Hadha and Hadhb protein levels remain elevated for at least 5 days following transfection with 1µg of mRNA containing equimolar amounts of *Hadha* and *Hadhb*. (C) Densitometry analysis of Hadha and Hadhb in panel B across three replicates. Hadha and Hadhb were first normalized to the Gapdh signal in each sample before normalization against the WT signal (set to 1). Dashed lines represent the average expression of Hadha and Hadhb in untreated TFP-D fibroblasts. d.p.t= days post-transfection.

To investigate the kinetics of Hadha and Hadhb expression following mRNA transfection, cellular lysates were collected at multiple timepoints after transfection and analyzed by western blot (Fig 4B). Expression of *Hadhb* peaked at 24 hours post-transfection, while the peak of *Hadha* expression was delayed to 48 hours post-transfection. The expression of each subunit remained significantly above baseline until day 5 with non-significant increases detectable on day 7. The calculated half-lives for Hadha and Hadhb protein were 3.7 and 2.8 days respectively. Given the two mRNAs were transfected in equimolar amounts and shared the same synthetic 5’ and 3’ UTR elements, the weaker, delayed recovery of Hadha protein may reflect a dependence on the expression of Hadhb.

### Intravenous administration of LNP-Hadha/Hadhb mRNA corrects metabolic deficits in aged TFP-D mice

Based on the robust *in vitro* response to Hadha/Hadhb mRNA transfection, therapeutic LNP-Hadha/Hadhb was evaluated in adult Hadhb^M404K/M404K^ mice (Fig. 5A). Mice received two doses of LNP at 1mg/kg via intravenous tail vein injection at 26 and 27 weeks of age. Muscle function and cold tolerance tests were assessed 24 and 48 hours after the second injection, respectively. Following the cold challenge, mice were euthanized and tissues were collected for biochemical and molecular analyses.

**Figure 5.**
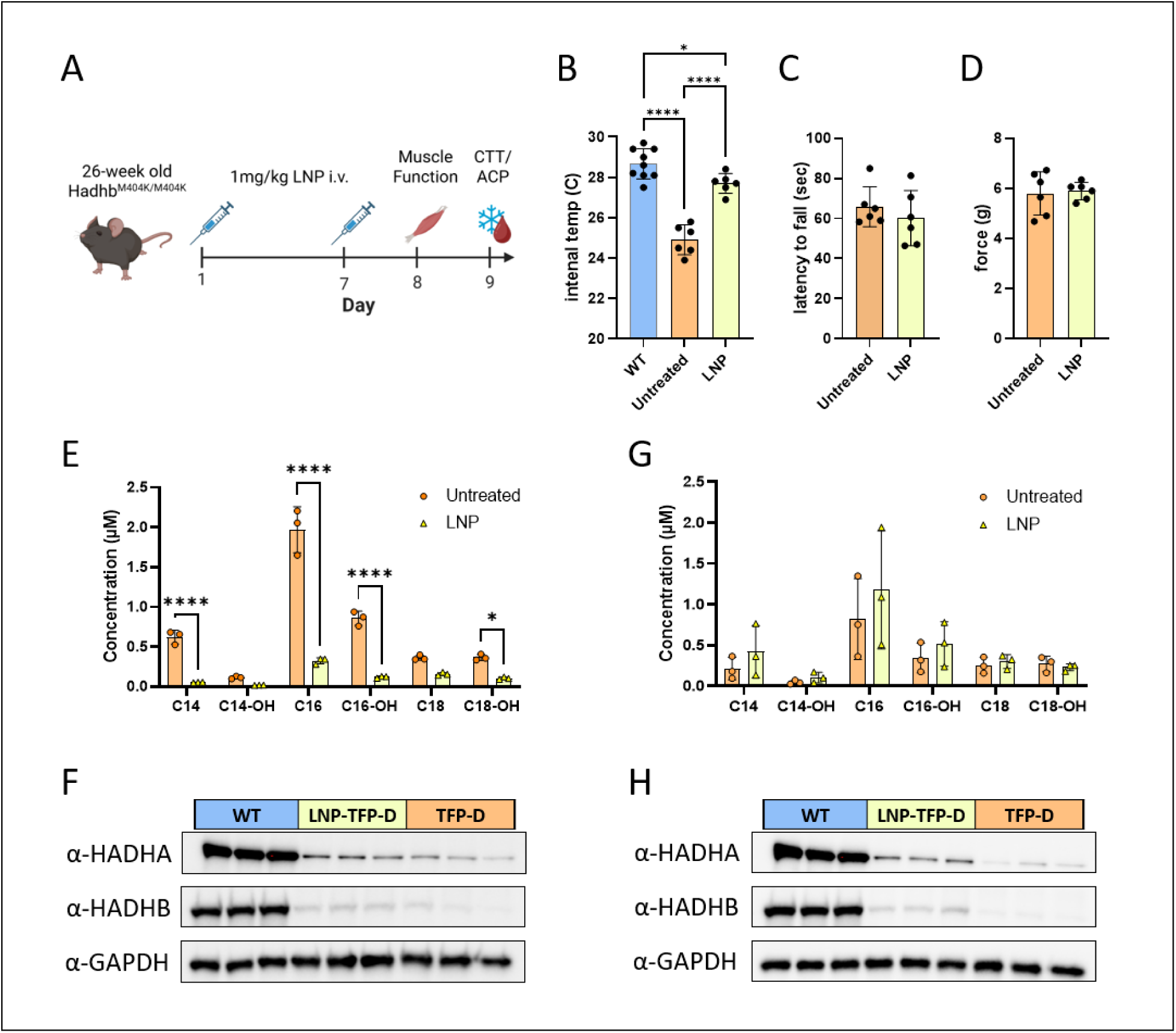
TFP-D mice show improved metabolism, biochemical correction, and partial protein restoration following 2 doses of LNP-Hadha/Hadhb. (A) Experimental schematic showing treatment and functional assessment timeline. (B) Cold tolerance testing demonstrates improved thermoregulation in LNP treated TFP D mice. One-way ANOVA with Tukey’s multiple comparison’s test. Data are presented as mean and standard deviation with individual mouse values shown. (C–D) Muscle function assessment by wire hang latency (C) and forelimb grip strength (D) shows no significant improvement following LNP Hadha/Hadhb mRNA treatment. Data are presented as mean and standard deviation with individual mouse values shown. Statistical non-significance determined by two-sided unpaired t-test. (E) Acylcarnitine profiling of male mice reveals a significant reduction in long chain acylcarnitine species in LNP treated mice. Two-way ANOVA with Šidák multiple comparisons test. (F) Representative immunoblot analysis of male liver demonstrates partial restoration of Hadha and Hadhb protein expression following LNP Hadha/Hadhb treatment. (G) Acylcarnitine profiling of female mice shows no significant reduction in long chain acylcarnitine species following treatment. Two-way ANOVA with Šidák multiple comparisons test. (H) Representative immunoblot analysis of female liver similarly shows partial restoration of Hadha and Hadhb protein expression. Gapdh is a loading control. All experimental groups contain equal numbers of male and female animals (12-week WT, n=8; 27-week TFP-D, n=6; 27-week TFP-D + LNP, n=6). *p<0.05, **p<0.01, ***p<0.001, ****p<0.0001

LNP-treated mice demonstrated significantly improved thermoregulation compared with age matched untreated TFP-D controls, but did not reach the level of wild-type mice (Fig. 5B). This assay is particularly relevant to TFP-D due to the requirement for long-chain fatty acid oxidation to support brown adipose tissue mediated thermogenesis and shivering associated energy production. Despite the rescued metabolic function, LNP-treated mice did not exhibit improvements in muscular endurance or neuromuscular function, as measured by the wire hang test (Fig. 5C), nor did they demonstrate increased forelimb grip strength (Fig. 5D).

Plasma acylcarnitine profiling revealed a significant reduction in the accumulation of multiple long-chain acylcarnitine species in LNP-treated male mice compared with untreated male controls (Fig. 5E). In contrast, treated female mice did not show a significant reduction in long-chain acylcarnitines (Fig. 5G). Despite these sex specific differences in biochemical correction, immunoblot analysis of liver tissue demonstrated partial restoration of both Hadha and Hadhb protein expression in LNP-treated male and female mice relative to untreated controls (Fig. 5F, H). Notably, both male and female LNP-treated mice exhibited improved thermoregulatory performance during cold exposure, indicating phenotypic improvement despite incomplete biochemical correction in female mice. Consistent with the muscle function findings, no increase in TFP protein expression was detected in skeletal muscle or heart tissue following treatment (Figure S4), suggesting limited LNP transduction in these tissues.

## Discussion

Mitochondrial trifunctional protein deficiency (TFP-D) remains associated with substantial morbidity and mortality despite newborn screening and strict dietary management, highlighting an urgent need for effective therapeutic interventions. Avoidance of fasting and supplementation with MCT oil or triheptanoin has provided a significant reduction in hospitalizations and hypoglycemia.^19^ While dietary strategies including triheptanoin can reduce the frequency and severity of rhabdomyolysis, they have limited benefits for exercise tolerance and cardiac function.^19–22^ In the open-label, long-term extension study, two of the 10 TFP-D patients on triheptanoin therapy died due to worsening cardiomyopathy.^19^ Dietary interventions rely on strict adherence, monitoring, and add a significant burden to patients and caregivers reducing quality of life. Here, we have identified a strategy for the stable treatment of TFP-D by periodic intravenous infusion of LNPs carrying a mixed cargo of *Hadha* and *Hadhb* mRNA.

Importantly, this strategy can overcome the dominant negative effect from endogenous Hadhb M404K protein in the mouse model. This factor will likely be key to the success of any genetic therapy for TFP-D, since it is expected that all patients living with TFP-D will produce some amount of variant HADHB that will competitively bind HADHA, as patients with total loss of function variants do not survive past the neonatal period. Given the shared pathophysiology of long-chain fatty acid oxidation disorders and the central role of the liver in generating circulating toxic intermediates, hepatic delivery of *Hadha*/*Hadhb* mRNA may likewise be sufficient to treat the related LCHAD-D caused by variants in *HADHA*.

Using patient relevant TFP-D fibroblasts derived from M404K mice, we show that restoration of fatty acid oxidation requires replacement of both *Hadha* and *Hadhb* mRNAs rather than augmentation of a single subunit. TFP-D fibroblasts exhibited marked accumulation of long chain acylcarnitine species, consistent with patient phenotypes. Transfection of TFP-D fibroblasts with *Hadhb* mRNA increased protein expression but failed to reduce long chain acylcarnitine accumulation. In contrast, dual delivery of *Hadha* and *Hadhb* mRNA reduced medium and long chain acylcarnitine species to near WT levels and increased protein expression. These findings reinforce results from an earlier plasmid-based study demonstrating that dual subunit replacement is required to restore TFP function in TFP-D fibroblasts.^14^ Treatment of affected fibroblasts with a *Hadhb* siRNA had the same rescue observed with dual LNP-*Hadha*/*Hadhb* delivery. By suppressing endogenous variant M404K Hadhb, the delivered WT Hadhb was able to form healthy TFP complexes without competition. The effectiveness of suppressing endogenous Hadhb strongly suggests a dominant negative effect from the M404K variant.

Regarding the discordant effect of isolated *Hadha* or *Hadhb* mRNA transfection (Figure 1), one possible explanation is that the western blot provides a snapshot of expression on day 3, but acylcarnitine profiling integrates enzyme activity over time. Effectively, if transfected WT *Hadhb* mRNA must first be translated and then outcompete the endogenous M404K Hadhb for Hadha binding, then there will be a delay in the rescue of acylcarnitine intermediates. Likewise, transfection with *Hadha* mRNA alone immediately increases the number of binding partners for endogenous M404K Hadhb subunits across all 3 three days, but does not ultimately correct the rapid turn-over of TFP complexes. Lower endogenous *Hadha* expression relative to endogenous *Hadhb* may predispose the system to destabilization and subunit imbalance, reinforcing the need for dual subunit replacement.

Computational and evolutionary analyses suggest that the M404K mutation disrupts a highly conserved methionine aromatic bridge critical for Hadhb structural stability and TFP complex integrity. While the M404K variant has not been reported in human patients, a F398L variant that similarly disrupts this structural motif has been identified.^23^ These motifs are highly conserved, and prior studies of the Hadhb found the F398 residue to be conserved across species. Extending this analysis we found that the identity and position of the Y357-M404-F398 bridge is conserved in the orthologous trifunctional protein beta subunit of *Sphaeroforma arctica*, a single celled eukaryote whose last common ancestor with humans existed over 800 million years ago, underscoring the structural importance of this bridge.^24^

LNP delivered mRNA therapies are emerging as a transformative approach for the treatment of inherited metabolic diseases and are already being advanced for disorders such as propionic acidemia, methylmalonic acidemia, ornithine transcarbamylase deficiency, and other fatty acid oxidation disorders like very long-chain acyl-CoA dehydrogenase deficiency (VLCAD-D) and medium-chain acyl-CoA dehydrogenase deficiency (MCAD-D).^10, 11, 25–27^ Notably, despite citing the use of the same liver targeting Moderna LNP in both the VLCAD-D and MCAD-D studies, tissue-specific protein restoration differed substantially between disease models, with correction limited to the liver in VLCAD-D but extending to cardiac and skeletal muscle in MCAD-D. The basis for this apparent disease dependent tropism remains unclear since it has not been disclosed whether the lipid formulation varied between the two studies, but opens the possibility that factors beyond LNP formulation such as disease specific metabolic stress, tissue permeability, cellular uptake, or mRNA processing may influence extrahepatic expression.

Consistent with the VLCAD-D findings, *in vivo* delivery of LNP-*Hadha*/*Hadhb* mRNA in the TFP-D mouse resulted in significant physiological improvement and partial biochemical correction restricted to the liver. In TFP-D male mice, LNP treatment resulted in a marked reduction in nearly all plasma long chain acylcarnitines despite the absence of increased protein abundance in heart or skeletal muscle. Female TFP-D mice did not exhibit comparable reductions in plasma acylcarnitines but demonstrated equivalent functional improvement and increased protein abundance by western blotting. While our analysis of liver, heart, and quadriceps tissue did not identify substantial sex-dependent differences in LNP tropism and protein expression, it remains possible that sex-specific uptake in non-tested tissues or systemic improvement below the limit of detection by western blot drives the improved acylcarnitines observed in male mice. Alternatively, the lack of improvement in female mice may reflect biological differences in either hepatic or systemic long chain acylcarnitine metabolism that are not observed at baseline, but become apparent following isolated correction of liver metabolism.

Among the key limitations of this study is that *in vivo* effects were examined after only two doses of LNP-mRNA where the treatment durability, multi-dose safety, and effect on long-term complications could not be evaluated. Nevertheless, the ability of LNP delivered mRNA to overcome a dominant negative variant and provide both biochemical and physiological benefit is highly encouraging. While mRNA therapies are inherently transient, weekly administration could represent a viable treatment paradigm. Furthermore, LNP-mRNA therapy could serve as a bridge treatment during childhood until growth is complete, at which point AAV based strategies could provide long-term correction. In patients with milder forms of TFP-D who may not require continuous LNP-mRNA therapy, this therapeutic may also be an effective intervention to breakthrough episodes of metabolic decompensation. Together, these findings reinforce hepatic LNP mRNA delivery as a promising therapeutic strategy for TFP-D while highlighting how disease context and biological sex may differentially influence metabolic correction and therapeutic response.

## Methods

### Cell Culture

Male WT and TFP-D mouse fibroblasts were isolated from ear punch biopsies according to standard methods. Fibroblasts were maintained in DMEM (Thermo Fisher Scientific, Cat. no. 11995073, Waltham, MA) supplemented with 15% fetal bovine serum (Millipore Sigma, Cat. no. F4135-500ML, Burlington, MA) and 1X concentrations of penicillin/streptomycin (Thermo Fisher Scientific, Cat. no. 15140122, Waltham, MA) and non-essential amino acids (Thermo Fisher Scientific, Cat. no. 11140050, Waltham, MA). For all *in vitro* experiments, cells were seeded at 100,000 cells per well in 1mL of complete medium in 12-well plates (USA Scientific, Cat. no. CC7682-7512, Ocala, FL).

### Plasmids

The sequence for eGFP was amplified from the pmax-GFP plasmid (https://www.addgene.org/177825/) and cloned into the pT7 backbone (https://www.addgene.org/178113/). The WT Hadhb cDNA sequence was synthesized by Genscript with a C-terminal FLAG tag and cloned into the pT7 backbone. The WT *Hadhb* cDNA sequence was subsequently cloned into the pDGP01 backbone without the C-terminal FLAG tag. The WT *Hadha* cDNA sequence was amplified from a WT mouse cDNA library and cloned into the pDGP01 vector backbone. Plasmid sequences were verified by whole plasmid sequencing and the integrity of the pDGP01 encoded poly(A) tail was confirmed by restriction digest and gel electrophoresis.

### In vitro mRNA synthesis

The template for eGFP and WT *Hadhb* mRNA used in figure 1 was produced from plasmid pT7-eGFP and pT7-Hadhb by PCR according to the methods of Doman, et al.^28^ The template for WT Hadha used in figure 1 was produced by AflII (New England Biolabs, Cat. no. R0520S, Ipswich, MA) restriction digest of the parent pDGP01 plasmid and gel purification of the linearized template. All mRNAs used in figure 1 were produced in small-scale 20µL IVT reactions using 1µg of linear template DNA. Small-scale IVT was performed using the HiScribe T7 mRNA kit with CleanCap Reagent AG (New England Biolabs, Cat. no. E2080S, Ipswich, MA) and complete N1-methylpseudouridine (Thermo Fisher Scientific, Cat. no. FERR0491T1, Waltham, MA) replacement according to the manufacturer’s recommendations. Small-scale mRNA reactions were purified by DNase I digestion of template DNA and standard LiCl precipitation as detailed in the HiScribe T7 mRNA kit.

For Figure 4 and the in vivo studies, WT Hadha and Hadhb mRNA were produced from the pDGP01 *Hadha* and *Hadhb* plasmid vectors in a large-scale synthesis reaction. mRNA was prepared by in vitro transcription (IVT) from a linearized plasmid template containing a T7 promoter, custom 5′/3’ UTRs, a Uridine-depleted and codon-optimized *Hadha* or *Hadhb* CDS, with an encoded poly(A) tail. mRNA IVT was performed as described by our group previously.^29^ Briefly, Plasmid DNA was linearized and IVT was performed at 37 oC with Promega T7 Enzyme Express Mix, a standard rNTP mix (ATP, CTP, GTP), and N1-methylpseudouridine triphosphate (TriLink Biotechnologies, Cat. no. N-1081, San Diego, CA). Co-transcriptional capping was performed with CleanCap AG 3′OMe (TriLink Biotechnologies, Cat. no. N-7413, San Diego, CA). Following transcription, residual plasmid DNA was digested with DNase I (New England Biolabs, Cat. no. G8002, Ipswich, MA) and quenched with 0.5 M EDTA (Promega, Cat. no. V4231, Madison, WI). Crude IVT material was purified using a multi-step downstream workflow consisting of tangential flow filtration (TFF) with a 100 kDa mPES cassette (Repligen, Cat. no. PP100LP2L, Waltham, MA) for buffer exchange into HEPES/NaCl/EDTA. Multimodal chromatography was used to purify the mRNA to remove rNTPs and DNA fragments, followed by a second TFF polishing step into citrate buffer pH 6.4. RNA was sterile-filtered with a 0.2 µm PES filter (Cytiva, Supor membrane, Wilmington, DE), and stored at -80°C. For LNP encapsulation, 1.5 mg of purified *Hadha* and *Hadhb* mRNA mixed at 1:1 molar ratio was encapsulated by Acuitas Therapeutics in Vancouver, British Columbia (proprietary ionizable cationic lipid, PEG-lipid ALC-0159, 1,2-distearoyl-sn-glycero-3-phosphocholine, and cholesterol). mRNA:LNP were rapidly mixed, followed by buffer exchange, sterile filtration, and fill/finish in glass vials for storage until administration. Ionizable cationic lipid and LNP compositions are described in Patent Applications WO/2017/004143A1.^30^

Dicer-substrate siRNA (DsiRNA) targeting mouse *Hadhb*were purchased from IDT as a TriFECTa RNAi kit (Integrated DNA Technologies, Cat. no. mm.Ri.Hadhb.13, Newark, NJ). See supplemental Table 1 for target site sequences of each DsiRNA.

### In vitro mRNA transfection

For *in vitro* transfections, mRNA/DsiRNA was complexed with Lipofectamine MessengerMax (Thermo Fisher Scientific, Cat. no. LMRNA001, Waltham, MA) at a ratio of 1µg of RNA/1µL of reagent. To prepare the complexes, both the RNA and Lipofectamine reagent were pre-diluted at 1µg RNA/1µL of Lipofectamine into separate tubes containing 50µL of Opti-MEM (Thermo Fisher Scientific, Cat. no. 11058021, Waltham, MA) and incubated at room-temperature for 10 minutes. The two parts were then combined, mixed, and incubated at room-temperature for an additional 10 minutes. The complexes were then added dropwise onto plates followed by gentle agitation for 10 seconds to achieve even distribution of complexes within the medium. Plates were then centrifuged at 220 x *g* (1000 rpm) in a benchtop centrifuge for 30 minutes to achieve rapid deposition of suspended complexes onto the cell monolayer. Cells were then returned to the incubator at 37°C and 5% CO_2_ while being careful to avoid disturbing the deposited complexes. For western blot and immunofluorescence experiments, transfection medium was replaced after 24 hours with normal culture medium. For palmitic acid challenge experiments, transfection medium was replaced with palmitic acid supplemented MEM after 2 hours. Preliminary flow cytometry experiments with eGFP mRNA demonstrated that a 2-hour incubation was sufficient to achieve robust transfection. The standard 24-hour incubation resulted in 81.3% of fibroblasts expressing eGFP, while the 2-hour incubation was only marginally reduced to 74.7% of fibroblasts expressing eGFP (data not shown).

### Western blot

Total cell lysates from either fibroblast cultures or mouse tissues were extracted by lysis in RIPA buffer containing 1X concentrations of protease inhibitors (Thermo Fisher Scientific, Cat. no. 78429, Waltham, MA). Proteins were denatured for 5 minutes at 98°C in 2X Laemmli buffer (Bio-rad, Cat. no. 1610747, Hercules, CA) with 5% β-mercaptoethanol. 5-15µg of sample was loaded onto a 12%-Mini-PROTEAN TGX precast gel (Bio-Rad, Cat. no. 4561044 or 4561046) and run alongside Precision Plus Protein Dual Color Standard (Bio-Rad, Cat. no. 1610374, Hercules, CA). Gel electrophoresis was performed at 200V until the 25kDa marker reached the end of the gel. Proteins were then transferred to a 0.4µm nitrocellulose membrane (Bio-rad, Cat. no. 1620115, Hercules, CA) via semi-dry transfer performed at 25V for 30 minutes at room temperature. Membranes were then blocked for 1 hour in 1X PBST + 1% nonfat dry milk (Bio-Rad, Cat. no. 170-6404, Hercules, CA). Membranes were then cut between the 37/50 kDa and 50/75 kDa markers for staining of the Gapdh (36kDa), Hadhb (50kDa), and Hadha (80kDa) bands with mouse α-GAPDH (Proteintech, Cat. no. 60004-1, Rosemont, IL), mouse α-HADHB (Proteintech, Cat. no. 67967-1, Rosemont, IL), and rabbit α-HADHA (ABclonal, Cat. no. A23892, Woburn, MA) respectively. Gapdh, Hadhb, and Hadha were stained for using primary antibodies at a 1:1,000 dilution in 1X PBST+1% BSA overnight at 4°C. Unbound primary antibody was removed by 3 washes with 1X PBST for 5 minutes each. Blots were then incubated for 1 hour at room temperature with the respective HRP conjugated Goat α-mouse Ig (SouthernBiotech, Cat. no. 1030-05, Birmingham, AL) or HRP conjugated Goat α-rabbit Ig (Bio-Rad, Cat. no. 403005, Hercules, CA) secondary antibodies diluted 1:10,000 in 1X PBST + 1% BSA. Unbound secondary antibody was removed by 3 washes with 1X PBST for 15 minutes each. Protein bands were visualized by chemiluminescence detection using the SuperSignal West Dura Substrate (Thermo Fisher Scientific, Cat. no. 34075, Waltham, MA) on a ChemiDoc imaging system (Bio-Rad, Hercules, CA). Image densitometry analyses were performed in ImageJ v1.54p.^31^

### Preparation palmitic acid MEM

To prepare the palmitic acid challenge medium, a normal minimal culture medium was first prepared containing MEM with 1X L-glutamine (Thermo Fisher Scientific, Cat. no. 11095-080, Waltham, MA) and supplemented with 10% FBS and 1X penicillin/streptomycin. A 200 mM (1000X) stock solution of palmitic acid (Millipore Sigma, Cat. no. P5585, Burlington, MA) in ethanol and a 400 mM (1000X) stock solution of L-Carnitine inner salt (Millipore Sigma, Cat. no. C0158, Burlington, MA) in water were then prepared followed by 0.2µm filter sterilization. To prepare the final medium, 175mg of fatty-acid free BSA (faf-BSA) (Millipore Sigma, Cat. no. A8806, Burlington, MA) was dissolved in 17.5mL of minimal culture medium. 50µL of palmitic acid stock solution was transferred to a new tube and the ethanol allowed to evaporate in a sterile biosafety cabinet. The faf-BSA containing medium was then added to the tube containing the dry palmitic acid residue followed by brief sonication and rocking at 37°C overnight to allow for saturation of the faf-BSA with palmitic acid. On the following day, the culture medium was supplemented with 50µL of L-carnitine stock solution and diluted to a final volume of 50mL with an additional 32.5mL of minimal culture medium followed by 0.2µm filter sterilization. The prepared media is stable for up to 3 months at 4°C.

### In vitro palmitic acid challenge

For the *in vitro* challenge experiments, normal culture or transfection medium was removed from the cells grown in 12 well plates (100,000 cells/well), and replaced with 420µL of pre-warmed palmitic acid challenge medium. Conditioned medium was collected after a further 72-hour incubation and stored frozen at -20°C until analysis.

### RT-qPCR of Hadha and Hadhb

Total RNA was extracted from cellular lysates of wild-type (WT) and affected fibroblasts using the RNeasy Mini Kit (Qiagen, Cat. no. 74104, Venlo, Netherlands) according to the manufacturer’s instructions with on-column DNase treatment (Qiagen, Cat. no. 79254, Venlo, Netherlands). RNA concentration and purity were assessed by 260nm absorbance. Complementary DNA (cDNA) was synthesized from extracted RNA using the High Capacity cDNA Reverse Transcription Kit (Thermo Fisher Scientific, Cat. no. 4368814, Waltham, MA) with oligo(dT)20 primers (Thermo Fisher Scientific, Cat. no. 18418020, Waltham, MA). Resulting cDNA was diluted to a final concentration of 25 ng per reaction for quantitative PCR analysis.

Quantitative real-time PCR (RT-qPCR) was performed using SYBR Green detection chemistry (Bio-Rad, Cat. no. 1725121, Hercules, CA) on a LightCycler 480 thermocycler (Roche, Basel, Switzerland). Three independent biological replicates were analyzed for both WT and affected fibroblasts, with three technical replicates performed per sample.

Primers for *Hadha* and *Hadhb* were designed to span exon-exon junctions, screened to avoid primer dimer formation, and validated to exhibit amplification efficiencies close to 100%. Primer sequences are available in supplemental table 2.

Absolute quantification of transcript copy number was performed using standard curves generated from serial dilutions of purified plasmids corresponding to each target gene. Standard curves were included on every qPCR plate and used to calculate absolute transcript copy numbers based on cycle threshold (Ct) values. Thermal cycling was performed according to the SYBR Green manufacturer’s recommendations, followed by melt curve analysis to confirm amplification specificity.

Transcript copy numbers for *Hadha* and *Hadhb* were calculated from their respective standard curves and normalized to input cDNA quantity.

### Quantification of acylcarnitines in cell medium

Acylcarnitines in the cell medium were analyzed as butyl esters using stable isotope dilution electrospray ionization-tandem mass spectrometry (ESI-MS/MS). A fixed (20 µL) amount of deuterium labeled acylcarnitine internal standards (IS) (d3-C0, d3-C2, d3-C3, d3-C4, d3-C8, d3-C14, and d3-C16) was mixed with 100 µL of cell medium in a 1.5 mL microcentrifuge tube. 800 µL of methanol was added to each sample to precipitate protein by vortex and centrifuging. The supernatant was transferred in an injection vial and dried under nitrogen, then esterified by heating with 3M butanolic HCl at 65 °C for 15 min. The butylated extract was dried under nitrogen and reconstituted in methanol:H2O (85:15, v:v). The samples (5 µL) were injected through a precolumn filter using ACQUITY I-Class UPLC system on Xevo TQ-S micro mass spectrometer (Waters Corporation, Milford, MA). The butylated acylcarnitines were analyzed using precursor ion scan of m/z 85. The acylcarnitine concentrations were calculated as the products of the peak intensity ratios of analytes to their correspondent IS multiplied by the response factors, which were the inverses of the slopes of calibration curves.

### Immunofluorescence

Fibroblasts were seeded at a density of 100,000 cells per well into 12-well plates with a 1cm glass coverslip in each well. After 24 hours, experimental wells were transfected with *Hadha* and/or *Hadhb* mRNA following the standard protocol and incubated for a further 48 hours at 37°C and 5% CO2. Cells were then fixed by a 10-minute incubation at room temperature with 4% paraformaldehyde dissolved in 1X PBS. Paraformaldehyde was then removed by 3 washes with 1X PBS followed by permeabilization for 5 minutes with 0.2% Triton X-100 in 1X PBS. Triton was then removed by 3 washes with 1X PBS followed by 1 hour of blocking at room temperature with 1X PBS + 5% normal goat serum (Thermo Fisher Scientific, Cat. no. 16210-064, Waltham, MA). Coverslips were then stained overnight at 4°C with 1:50 mouse α-HADHB and 1:200 rabbit α-HADHA diluted in 1X PBS + 1% BSA. Unbound primary was then removed by 3 washes for 5 minutes each in 1X PBS. Coverslips were then stained with Goat α-mouse Ig AF488 (Thermo Fisher Scientific, Cat. no. A11017, Waltham, MA) and Goat α-rabbit Ig AF596 (Thermo Fisher Scientific, Cat. no. A11071, Waltham, MA) fluorescent secondary antibodies diluted 1:1,000 in 1X PBS + 1% BSA for 1 hour at 37°C. Coverslips were then counterstained with 0.5X concentration DAPI diluted in 1X PBS for 2 minutes. Excess secondary and DAPI was then removed by 3 washes in 1X PBS for 15 minutes each. Coverslips were then mounted onto slides with 100% glycerol, kept shielded from light, and imaged within 30 minutes of mounting.

### Computational Protein Stability Analysis

#### The PAStE Algorithm

The PAStE algorithm^32^ was used as implemented in OSPREY 3.3^15^ to compute thermodynamic stability changes (ΔΔG) upon mutation. PAStE computes provably accurate

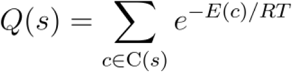

approximations of the partition function for each protein sequence by enumerating conformations in order of increasing energy using A* search and accumulating their Boltzmann-weighted contributions.

For each sequence s, the partition function is defined as:

where the sum is over all conformations c in the conformational space C(s) of sequence s, E(c) is the energy of conformation c, R is the gas constant, and T is the temperature (298.15 K). The ΔΔG between a mutant sequence and wild-type is then:

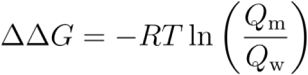

where *Q_m_* is the partition function of the mutant and *Q_w_* is the partition function of the wild-type sequence.

A key of PAStE is its use of ε-approximation to provide guaranteed bounds on the computed ΔΔG. It maintains rigorous upper and lower bounds on each partition function throughout the computation, ensuring that the reported ΔΔG values are accurate to within the specified tolerance. PAStE employs a gradient-descent algorithm to iteratively tighten upper and lower energy bounds, thereby refining the corresponding partition function bounds. A window criterion allows early termination when the bounds of the mutant and wild-type partition functions no longer overlap, establishing the sign of ΔΔG with certainty.

#### Computational Setup

The input structure was the crystal structure of the human mitochondrial trifunctional protein (PDB ID: 6DV2). The structure was preprocessed by stripping non-protein atoms and adding hydrogen atoms. Custom residue templates were generated for Nε-acetyl-lysine (ALY, template ID: K03) and Nε-succinyl-lysine (SLL, template ID: K04), including template geometries, atomic coordinates, and rotamer libraries. OSPREY employs an energy function combining AMBER parm96 force field parameters^33^ with atomic partial charges from the ff94 parameter set,^34^ together with the EEF1 implicit solvation model.^35^ For handling non-proteinogenic amino acids, the force field parameterization follows the methodology described in Wang, 2021.^36^

The conformational space included residue 404 (the mutation site; PDB residue B781) with wild-type, ALY, and SLL rotamers, and eight neighboring residues (T357, Y358, A359, W398, F399, A400, Y403, G405; PDB residues B734–B736, B775–B777, B780, B782) with wild-type rotamers. All flexible residues used continuous rotamer optimization. PAStE was configured with ε = 0.1, a maximum of 50,000 conformations per partition function evaluation, and the window overlap criterion enabled. Energy calculations used 20 CPU cores.

### Mouse husbandry

All animal experiments were conducted in accordance with the guidelines of the Institutional Animal Care and Use Committee (IACUC) at Duke University and approved under protocol A004-23-01. TFP-D mice have been previously described.^13^ All mice were maintained under standard specific-pathogen-free conditions with a 12-hour light/dark cycle and ad libitum access to standard mouse chow (LabDiet, Cat. no. 3005740-220, St. Louis, MO) and water. Homozygous affected not treated and affected treated mice were analyzed from 26-27 weeks of age. Healthy WT controls were analyzed at 12 weeks of age. Each group had 3 male and 3 female mice.

Mice were euthanized according to approved humane protocols, and tissues were collected immediately, frozen on dry ice, and stored at −80 °C until processing. Tissue samples were subsequently used for protein extraction for western blotting and plasma acylcarnitines were assessed.

### LNP administration in mice

Homozygous affected TFP-D mice were treated at 26 and 27 weeks of age with LNP-*Hadha*/*Hadhb* at a dose of 1 mg/kg via intravenous tail vein injection. Prior to administration, LNP-*Hadha*/*Hadhb* was thawed on ice and diluted 1:5 in Dulbecco’s phosphate-buffered saline (Thermo Fisher Scientific, Cat. no. 14200075, Waltham, MA). Injections were performed via the right lateral caudal vein at week 26 and the left lateral caudal vein at week 27 to minimize injection-site trauma and scar tissue formation.

Affected, untreated control TFP-D mice received vehicle control injections consisting of DPBS alone and were handled identically to the treatment group.

### Cold tolerance test

Forty-eight hours after the final LNP treatment, mice were transferred to a cold room maintained at 4°C with ad libitum access to food and water. Mice were housed in groups of three according to treatment regimen for the duration of the assay. Group housing was maintained consistently across all experimental groups to control for potential effects of social thermoregulation. Animals were monitored continuously and remained in the cold environment for a maximum of 4 hours or until core body temperature reached 25°C, at which point the experiment was terminated to prevent hypothermia.

Core body temperature was measured rectally using a Thermalert Th-5 digital temperature probe (Physitemp, Clifton, NJ) connected to a temperature monitoring system. Temperature measurements were recorded at baseline (0 hours) and at 1-hour intervals thereafter until the end of the experiment (4 hours or when body temperature reached ≤25 °C).

### Muscle Function Assessment

Mouse muscle function was assessed twenty-four hours after the final LNP treatment (week 27; post-treatment). All testing was performed during the light cycle and mice were allowed to acclimate to the testing room prior to assessment.

### Wire Hang Test

Muscular endurance and neuromuscular function were evaluated using the wire hang test. Mice were placed on a wire grid, which was then gently inverted to position the animals upside down, and latency to fall was recorded. Each mouse performed three wire hang trials with a 5-minute rest period between trials. Latency values from the three trials were averaged for each mouse prior to data analysis to account for intra-animal variability. Following completion of the wire hang test, mice were allowed a 10-minute rest period before grip strength testing.

### Grip Strength Test

Forelimb grip strength was assessed using a grip strength meter equipped with a DFE II force transducer (Chatillon, Jamaica, NY). Only the forelimbs were allowed to grasp the force sensor, and mice were gently pulled backward in a horizontal plane until grip was released. The peak force generated during each pull was recorded. Each mouse underwent three consecutive pulls followed by a 5-minute rest period, and this sequence was repeated three times for a total of three grip strength attempts per mouse. Grip strength values were averaged prior to analysis.

During both wire hang and grip strength testing, mice had continuous access to food and water.

## Software and Data Availability

OSPREY is freely available at https://github.com/donaldlab/OSPREY3. The input files, custom residue templates, and analysis scripts used in this study are available. The datasets generated and analyzed during the present study are available from the corresponding author upon reasonable request. Additional information required to interpret, verify, or extend these findings is also available from the corresponding author upon reasonable request.

## Acknowledgements

This work was supported by Duke Department of Pediatrics Research Funding (Dwight D. Koeberl) and the National Institute of General Medical Sciences (R35 GM-144042 to Bruce R. Donald).

## Author Contributions

Conceptualization, T.V.B. and D.D.K.; methodology, T.V.B., T.T., B.F.M, S.P.Y., B.R.D., and D.D.K.; investigation, T.V.B., T.T., H.Z., S.L. (in vitro/in vivo), B.F.M (RNA synthesis/encapsulation), and L.Z. (in silico); data curation and visualization, T.V.B, T.T., and L.Z.; supervision, S.P.Y., B.R.D, M.L, C.T., D.D.K.; writing – original draft, T.V.B., T.T., H.Z., B.F.M, L.Z., and B.R.D.; writing – review and editing, T.V.B and D.D.K.

