## Supplemental Information for "Dual mRNA delivery corrects physical and biochemical deficits in a mouse model of trifunctional protein deficiency"

**Supplemental Tables**

**Table S1. siRNA target sequences and catalog numbers.**

| **Name** | **Target Sequence** | **Cat. no.** |
|---|---|---|
| mm.Ri.Hadhb.13.1 | aggttttcgataatcacagcttctg | N/A |
| mm.Ri.Hadhb.13.2 | aggatgttgttgattatatcatctt | N/A |
| mm.Ri.Hadhb.13.3 | gtccttactcagtaaattcagattg | N/A |
| Universal Negative Control DsiRNA | N/A | 51-01-14-03 |

**Table S2. RT-qPCR Primers**

| **Name** | **Sequence** |
|---|---|
| Hadha Forward | AACGTATGCCATGACTGTCCC |
| Hadha Reverse | TCCAGCCTTCACAGCATCAATTA |
| Hadhb Forward | CCAAAAGTTTTAGAAAAGGCAGGG |
| Hadhb Reverse | GCTAAAATCTGGCCTGAGAAGG |

**Supplemental Figures**

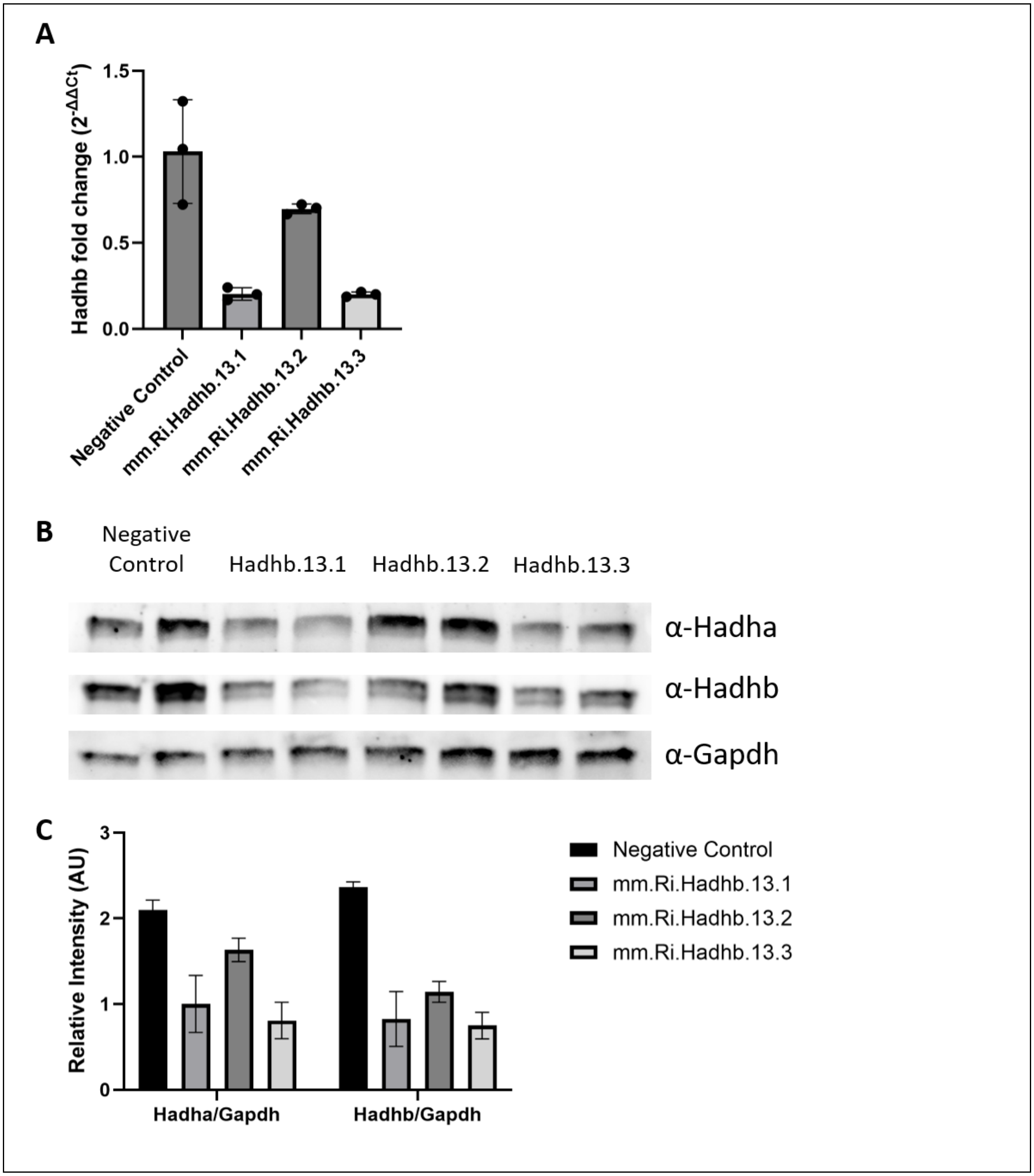


**Figure S1. Screening of Hadhb targeting siRNAs.** WT mouse fibroblasts were transfected with 1µg (A) or 0.5µg (B-C) of siRNA targeting *Hadhb* or universal negative control DsiRNA using the standard transfection protocol with Lipofectamine MessengerMax. RNA and protein lysates were collected 72 hours after transfection and used for relative mRNA and protein quantification. (A) RT-qPCR results for *Hadhb* mRNA relative to *Actb* (β-actin) mRNA. (B) WB results for Hadha, Hadhb, and Gapdh. (C) Image densitometry quantification of Hadha and Hadhb protein relative to Gapdh shown in (B).

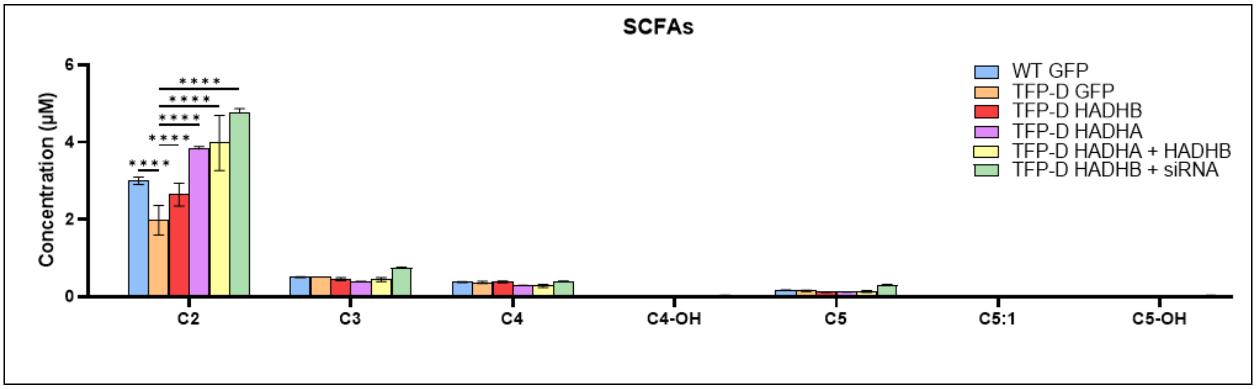


**Figure S2. Short chain acylcarnitine profile for samples in Figure 1D and E.** Data are presented as mean ± standard deviation from three independent experiments (n = 3). Statistical significance was determined by two-way ANOVA with Dunnett's multiple comparisons between all samples and 2µg GFP treated TFP-D. ****p<0.0001.

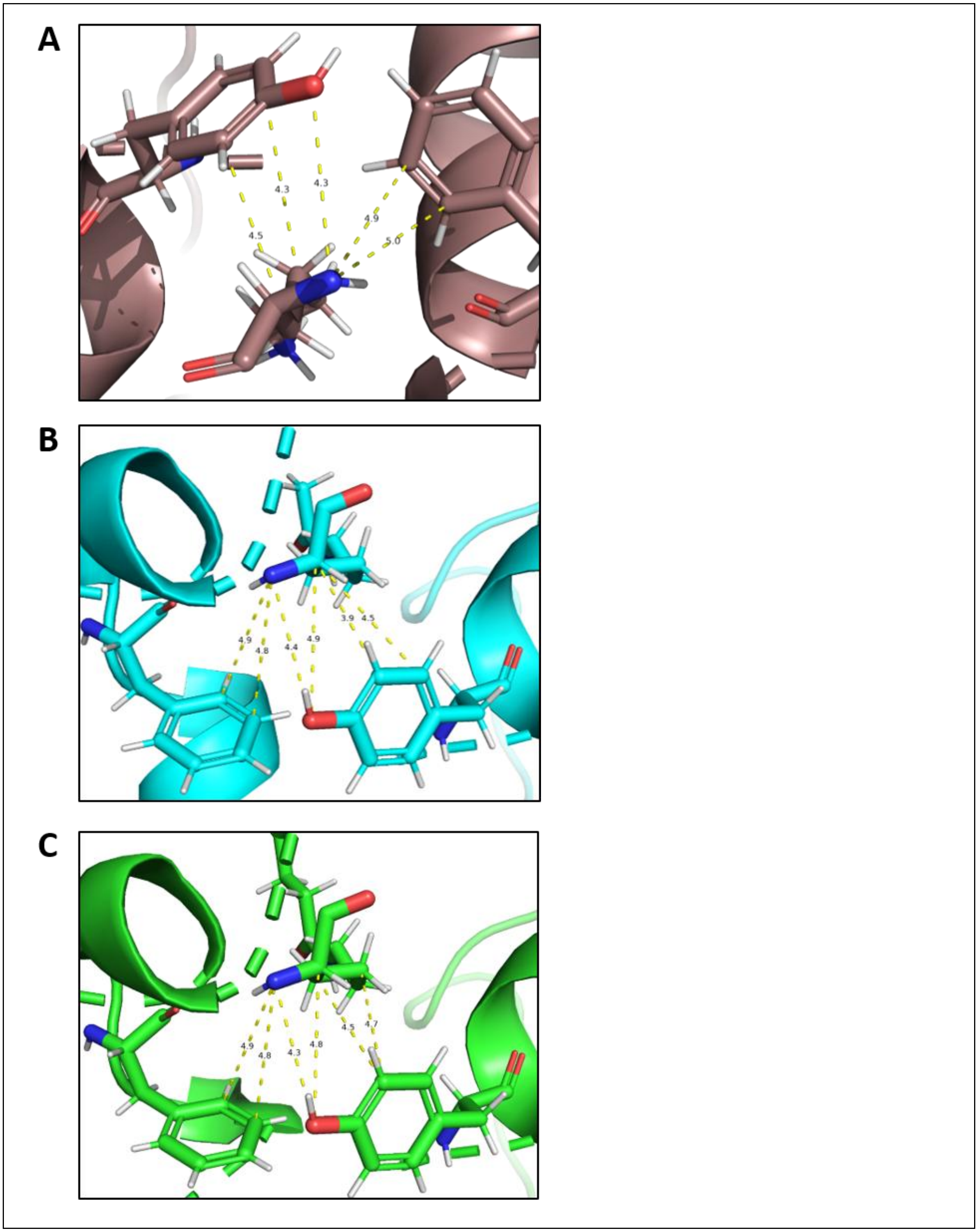


**Figure S3. Structural models of lysine variants at position 404 of Hadhb (PDB:6DV2).** (A) The unmodified M404K side-chain is depicted in the context of the surrounding methionine-aromatic bridge residues Y358 and F399 (shown in stick representation). (B) The M404K-ALY variant acetyl-lysine side chain (shown in stick representation) is depicted in the context of the surrounding methionine-aromatic bridge residues Y358 and F399 (shown in stick representation). The acetyl group extends the lysine side chain beyond the volume occupied by the native methionine, resulting in steric clashes with the aromatic cage formed by Y358 and F399. (C) The M404K-SLL variant succinyl-lysine side chain (shown in stick representation) is depicted in the context of the surrounding methionine-aromatic bridge residues Y358 and F399 (shown in stick representation). The bulkier succinyl group introduces greater steric strain than the acetyl modification, consistent with the larger predicted destabilization. The carboxylate moiety of the succinyl group may also introduce unfavorable electrostatic interactions within the hydrophobic aromatic cage. Key interatomic distances (in Å) are annotated.

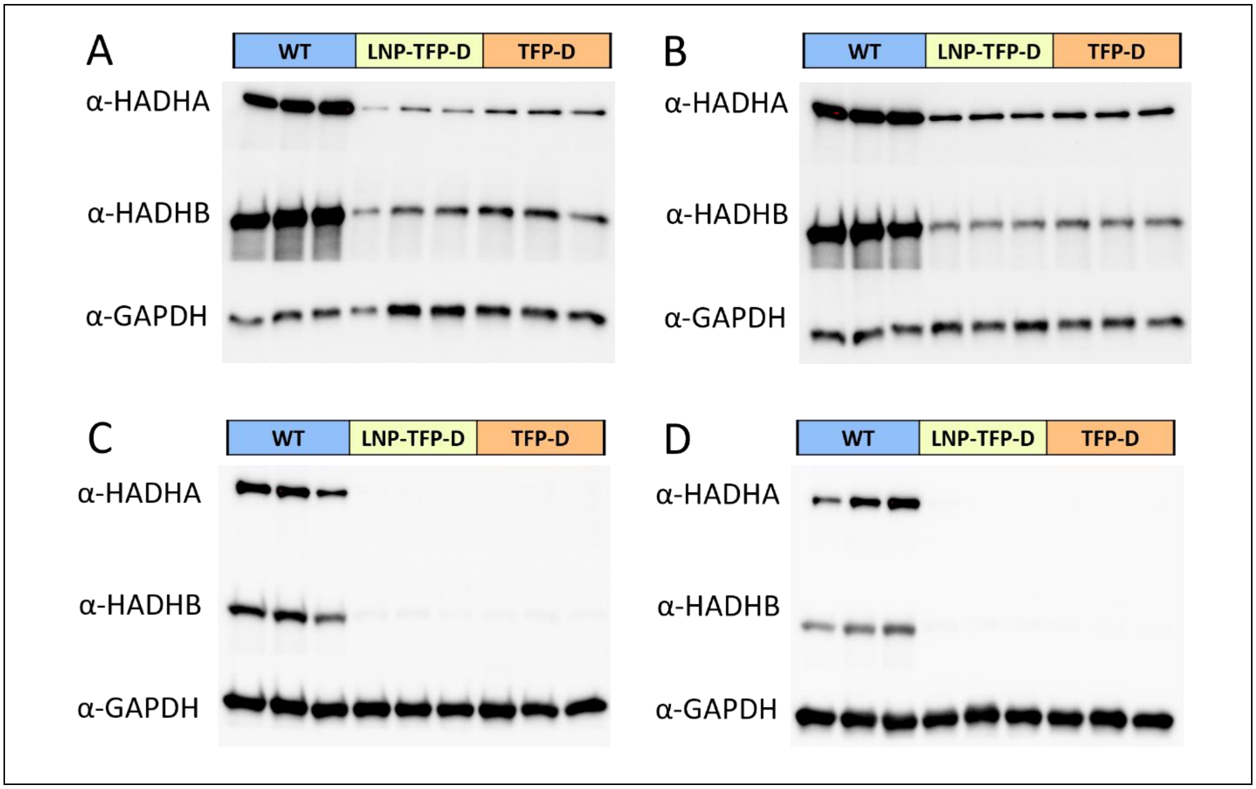


**Figure S4. Representative immunoblots of heart and quadriceps tissue.** (A-B) Expression of TFP subunits in heart tissue from male (A) and female (B) mice. (C-D) Expression of TFP subunits in quadriceps tissue from male (C) and female (D) mice.
